# Transcription-directed membrane association organizes the chloroplast nucleoid structure

**DOI:** 10.1101/2023.05.12.540520

**Authors:** V. Miguel Palomar, Sho Fujii, M. Hafiz Rothi, Sarah Jaksich, Adriana N. Coke, Joyful Wang, Andrzej T. Wierzbicki

## Abstract

DNA is organized into chromatin-like structures, which support the maintenance and regulation of genomes. A unique and poorly understood form of DNA packaging exists in chloroplasts, which are endosymbiotic organelles responsible for photosynthesis. Chloroplast genomes, together with associated proteins, form membraneless structures known as nucleoids. The internal arrangement of the nucleoid, molecular mechanisms of DNA packaging, and connections between nucleoid structure and gene expression remain mostly unknown. We show that *Arabidopsis thaliana* chloroplast nucleoids have a unique organization driven by DNA binding to the thylakoid membranes. DNA associated with the membranes has high protein occupancy, reduced DNA accessibility, and is highly transcribed. In contrast, genes with low levels of transcription are further away from the membranes, have lower protein occupancy, and higher DNA accessibility. Disruption of transcription at specific genes in sigma factor mutants causes a corresponding reduction in membrane association, indicating that RNA polymerase activity causes DNA tethering to the membranes. We propose that transcription organizes the chloroplast nucleoid into a transcriptionally active membrane-associated core and a less active periphery.

## INTRODUCTION

Packaging of DNA with proteins and RNAs is essential for genome maintenance and regulation. In the eukaryotic nucleus, chromatin is a complex multilevel structure, which supports many aspects of genome function. However, canonical eukaryotic chromatin is not the only form of DNA packaging. Alternative modes of DNA organization are present in mammalian sperm cells ^1^, dinoflagellates ^2^, archaea ^3^, bacteria ^4^, and viruses ^5^. Structural arrangements of DNA in those tissues or organisms relies on distinct protein machineries and provide unique functional impacts.

Mitochondria and plastids originated from bacterial ancestors and contain their own DNA. This DNA is packaged into chromatin-like structures known as nucleoids. Organellar genomes have special properties that have developed over more than a billion years of co-evolution with the eukaryotic cell host. The DNA packaging mechanisms of mitochondria and plastids are clearly distinct from what is known in either eukaryotic nuclei or bacteria, making organellar nucleoids an especially unique cases of DNA packaging ^6, 7^. In line with this, nucleoid-associated proteins (NAPs) in the chloroplasts of land plants are atypical. NAPs are thought to help organize DNA and support transcription ^8^. However, chloroplasts of land plants do not contain typical bacterial NAPs like HU or IHF. Also, chloroplast NAPs have little in common with nuclear chromatin proteins, and their biochemical functions remain mostly unknown ^9^. This indicates that mechanisms of DNA packaging in plastid nucleoids are likely distinct from their bacterial or nuclear counterparts.

The mechanisms of plastid DNA packaging, the internal structure of nucleoids, and their functional impacts remain poorly understood ^6^. In most land plants mature chloroplast nucleoids are localized in the stroma, near thylakoid membranes ^10, 11^. Electron microscopy studies found that the chloroplast nucleoids have a dense and protein-rich central body, which is resistant to high salt treatment and contains 30 to 50% of DNA. The remaining DNA may be observed as protruding fibrils with weaker protein binding ^12–14^. The *in vivo* relevance of the central body remains unclear due to the limitations of sample fixation for electron microscopy and insufficient resolution of light microscopy. The function of the central body is also unclear with some evidence favoring this structure as the site of active transcription ^11, 15^ while others suggesting that it may be associated with lower levels of transcription ^16^.

An important property of both bacterial and nuclear DNA packaging is the presence of sequence-specific structural features. This means that individual loci have unique patterns of protein binding, DNA accessibility, and subcellular or subnuclear localization to support the unique properties of each locus. In contrast, it is unknown if the chloroplast nucleoid includes a widespread presence of sequence-specific structural features. Some evidence supports the possibility of the entire genome adopting uniform structural properties ^17^, while other studies suggest some level of sequence-specificity ^18, 19^. It remains unknown if chloroplast nucleoids have a sequence-specific pattern of DNA accessibility, 3D genome organization or suborganellar localization. Therefore, it is difficult to predict if transcription, replication, and other processes involving DNA are supported by or control the arrangement of the chloroplast nucleoid.

To determine if the chloroplast nucleoid has a sequence-specific pattern of structural features, we adopted a broad range of genome-wide approaches. We found that the overall protein occupancy on DNA is dominated by transcriptional machinery and has an impact on restricting the accessibility of DNA. The chloroplast genome has a highly complex and sequence-specific pattern of association with thylakoid membranes. Membrane association is correlated with plastid encoded RNA polymerase (PEP) binding. Moreover, disruption of PEP transcription leads to disruptions of DNA membrane association, which indicates that transcription is required for bringing certain genes to the membranes.

## RESULTS

### Protein binding to chloroplast DNA is dominated by PEP

To test if the chloroplast nucleoid has a sequence-specific pattern of protein occupancy on DNA, we performed an *In vivo* Protein Occupancy Display (IPOD) assay ^20, 21^. This method relies on formaldehyde crosslinking, purification of protein-DNA complexes by phenol extraction, and high throughput sequencing. IPOD performed with purified Arabidopsis chloroplasts revealed a highly complex pattern of protein occupancy on DNA (Fig. 1A), which is consistent with prior low-resolution findings ^11, 19^. Its most obvious property is strong enrichment on highly transcribed genes, which is reminiscent of the previously reported pattern of Plastid Encoded RNA Polymerase (PEP) binding to DNA ^22^ (Fig. 1B). This result is also consistent with the known strong representation of RNA polymerase occupancy in bacterial IPOD occupancy traces^21^. Within individual genes, protein occupancy is often enriched on promoter regions, which are known to be preferentially bound by PEP ^22^ (Fig. 1C). Consistently, the IPOD signal and PEP binding to DNA are highly and significantly correlated on both annotated genes (Fig. 1D) and throughout the entire genome (Fig. S1A). As much as 82% of IPOD variance may be explained by PEP binding (Fig. 1D). The presence of a few genomic regions that do not follow this correlation may be explained by the presence of other nucleoid-associated proteins with weaker or transient binding ^8, 23–25^. Additionally, more proteins may bind DNA without sequence-specificity, as non sequence-specific interactions may be undetectable by IPOD. Together, these results indicate that there is a complex and sequence-specific pattern of protein binding to DNA in chloroplast nucleoids and that this binding is dominated by PEP.

**Figure 1.**
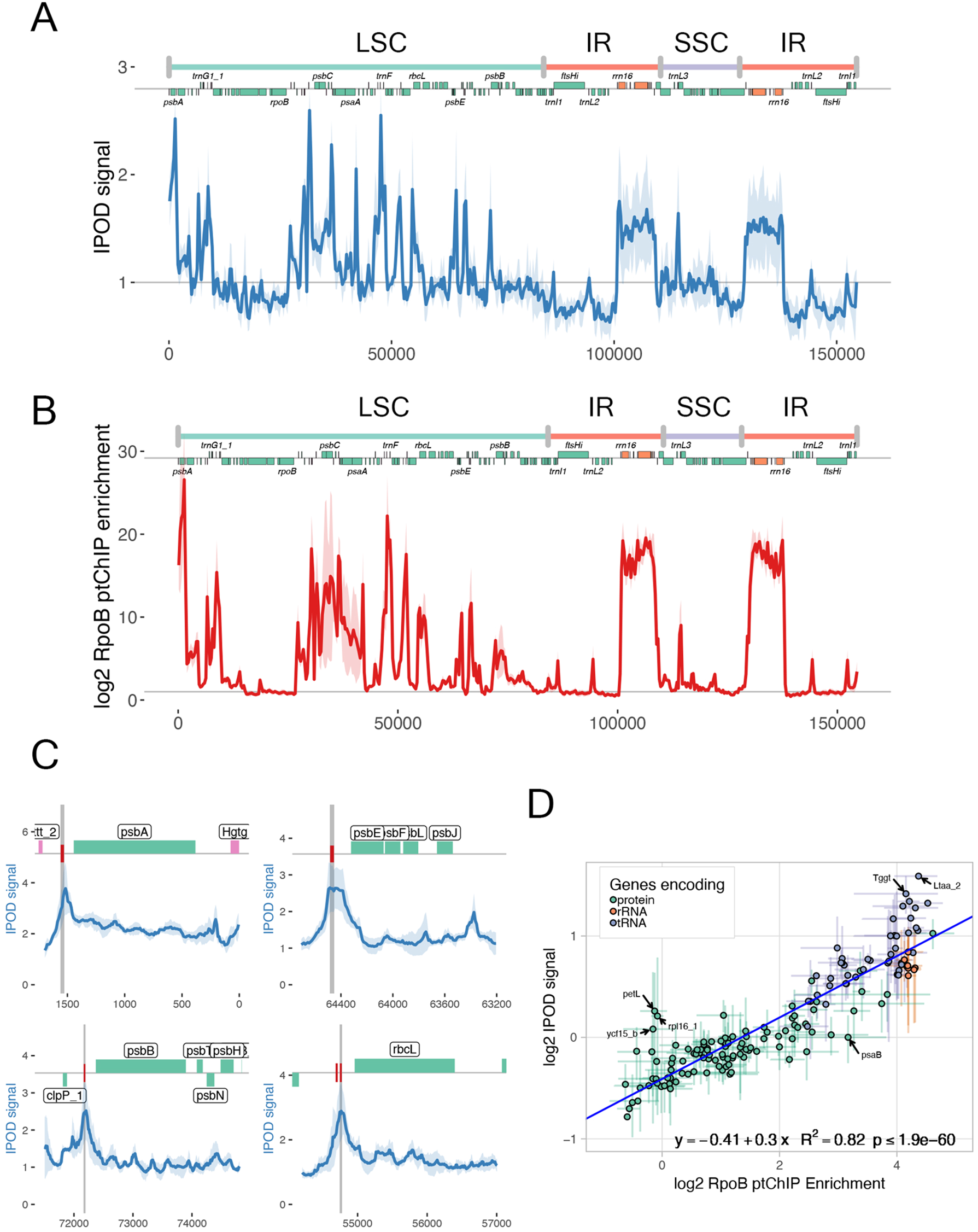
Protein binding to chloroplast DNA is dominated by PEP. A. Genome-wide pattern of protein occupancy on DNA detected by IPOD. Signal from IPOD in Col-0 wild-type plants was calculated in 50-bp genomic bins and plotted throughout the entire plastid genome. Genome annotation including genomic regions, positions of annotated genes ^22^, and names of selected individual genes is provided on top of the plot. Average enrichments from four independent biological replicates are shown. The light blue ribbon indicates standard deviation. B. Previously published genome-wide pattern of PEP binding to DNA ^22^. Signal enrichment from ptChIP-seq using αRpoB antibody in Col-0 wild-type plants was calculated in 50-bp genomic bins and plotted throughout the entire plastid genome. Average enrichments and standard deviation from three independent biological replicates are shown. C. Preferential protein occupancy on gene promoters. IPOD signal enrichment from Col-0 wild-type was calculated in 10-bp genomic bins and plotted at *psbA*, *psbE*, *psbB* and *rbcL* loci. Average enrichments from four independent biological replicates are shown. Light blue ribbons indicate standard deviations. Gray vertical line indicates positions of the annotated promoters. Genome annotation is shown on top. D. Protein occupancy and PEP binding are significantly correlated. IPOD signal and RpoB ptChIP-seq signal ^22^ were compared on annotated genes. Data points are color-coded by function and show averages from three (ptChIP-seq) or four (IPOD) biological replicates. Error bars indicate standard deviations. The blue line represents the linear regression model.

### PEP-occupied genes have reduced DNA accessibility

To test if protein occupancy is negatively correlated with DNA accessibility, we developed a modified version of the Assay for Transposase-Accessible Chromatin (ATAC-seq), which is a method used to study nuclear genomes and relies on fragmentation of the genome by engineered Tn5 transposomes ^26^. In this method, accessible DNA serves as a good substrate for transposon integration, but strong protein binding prevents transposon insertion. We optimized ATAC-seq to study plastid nucleoids and refer to this approach as ptATAC-seq. In our modified protocol, purified chloroplasts are crosslinked with formaldehyde as described previously ^22^, lysed with a hypotonic buffer, incubated with Tn5 transposomes, and assayed by high throughput sequencing. Purified (naked) DNA that has not been crosslinked is used as a control.

ptATAC-seq on wild type Arabidopsis chloroplasts revealed a relatively complex pattern of Tn5 insertions into the plastid genome (Fig 2A). Observed effects were low and variation within the eleven biological replicates of this experiment was high (Fig. S2A), which explains why only small subsets of the genome had significant enrichments or depletions of Tn5 integration (Fig. 2A). To test if low levels of Tn5 integration correspond to high PEP binding and overall protein occupancy, we split 50 bp genomic bins into groups with significant enrichment of Tn5 integration (accessible), significant depletion of Tn5 integration (protected) or no significant change (undetermined) (Fig. 2B). Genomic bins marked as accessible had low levels of PEP binding detected by RpoB ptChIP-seq, while bins marked as protected had high levels of PEP binding (Fig. 2C). This indicates that high levels of PEP binding are associated with depletion in Tn5 integration. This may be interpreted as evidence of at least partial DNA protection by PEP and associated proteins.

**Figure 2.**
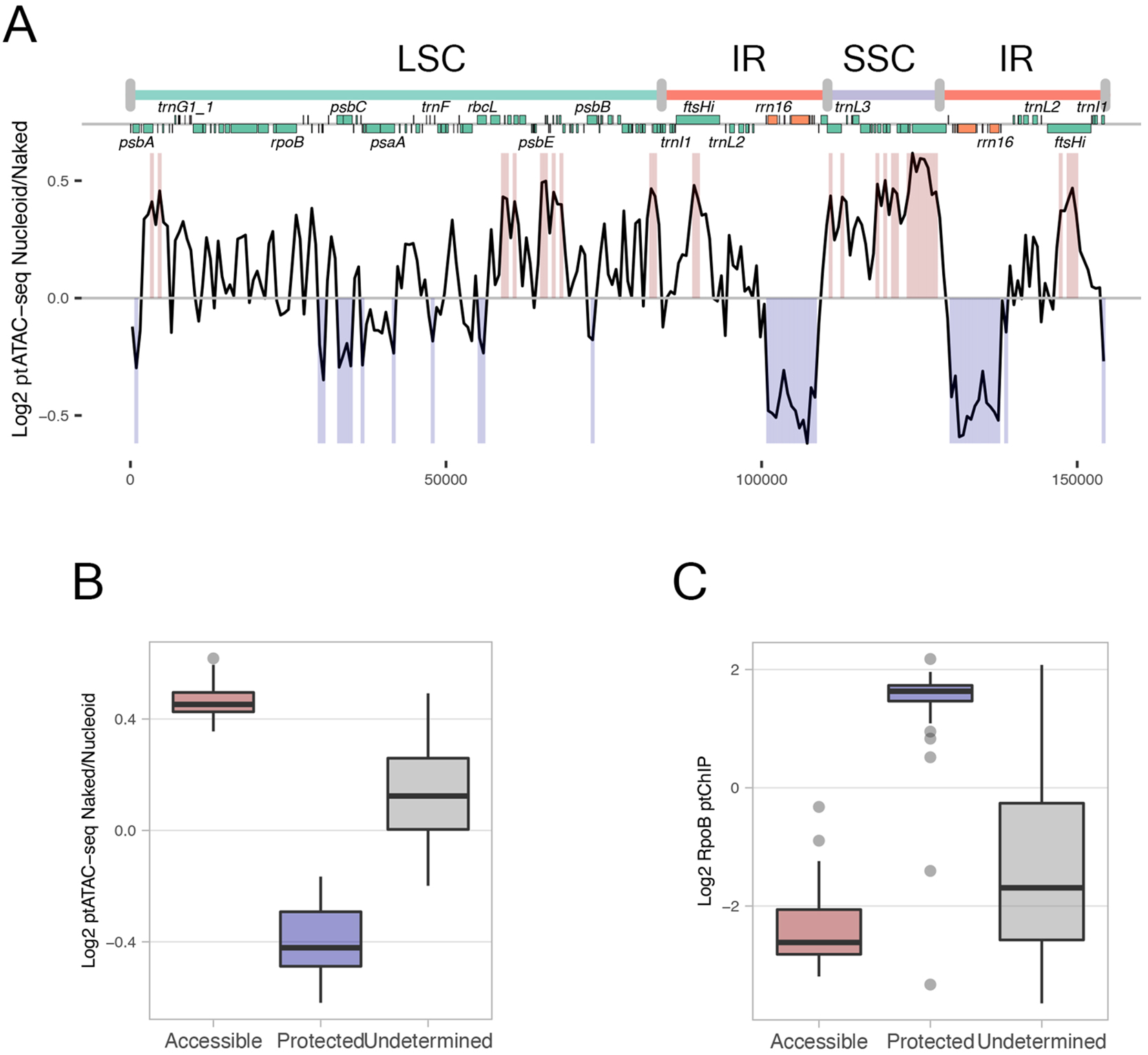
Transcribed genes have reduced DNA accessibility to Tn5 transposase. A. Genome-wide pattern of DNA accessibility detected by ptATAC-seq. Signal from ptATAC-seq in Col-0 wild-type plants was calculated in 50-bp genomic bins and plotted throughout the entire plastid genome. Y axis represents ratio of insertions into crosslinked nucleoid to insertions into purified (naked) DNA. Genome annotation including genomic regions, positions of annotated genes ^22^, and names of selected individual genes is provided on top of the plot. Average signal from eleven independent biological replicates is shown. Red shading indicates significant accessibility and blue shading indicates significant protection identified using a negative binomial model FDR < 0.05. Individual biological replicates are shown in Fig. S2A. B. Identification of genomic bins with significant accessibility or protection. Negative binomial model (FDR <= 0.05) was used to identify 50 bp genomic bins with significant enrichment (accessible) or depletion (protected) of Tn5 insertions. Regions with no significant change (FDR > 0.05) were identified as undetermined. Individual data points within boxplots are averages from eleven biological replicates. C. Protected genomic regions have high PEP binding. Previously published RpoB ptChIP-seq signal ^22^ plotted on genomic bins identified as accessible, protected or undetermined (Fig. 2B). Individual data points within boxplots are averages from three biological replicates.

### Chloroplast nucleoid is organized by association with the membranes

An important property of chloroplast nucleoids is their association with the thylakoid membranes ^10, 11^. DNA interactions with the membranes may involve a subset of genes in a sequence-specific pattern. This has been suggested by studies in spinach ^18, 27^. Interestingly, other studies suggest that DNA association with the membranes may have limited or no sequence specificity ^17^. To distinguish between these alternative scenarios, we developed an assay to study sequence-specific DNA-membrane associations on the genome-wide scale (Fig. 3A). The <u>Sol</u>uble-<u>In</u>soluble Nucleoid Fractionation <u>A</u>ssay (SOLINA) is based on a well-established approach to separate thylakoid membranes and stroma by centrifugation of lysed chloroplasts ^28–32^. In the SOLINA assay, purified Arabidopsis chloroplasts are crosslinked with formaldehyde and lysed using hypotonic buffer, then DNA is fragmented by partial digestion with Micrococcal nuclease (MNase). Subsequently, the sample is fractionated by centrifugation. The insoluble fraction (pellet) is expected to contain the membranes together with crosslinked DNA. The soluble fraction (supernatant) is expected to contain stroma and DNA fragments, that were not crosslinked to the membranes. DNA from both fractions is quantified by high throughput sequencing and the ratio of pellet to supernatant signals is interpreted as enrichment of a particular sequence in the membranes (Fig. 3A).

**Figure 3.**
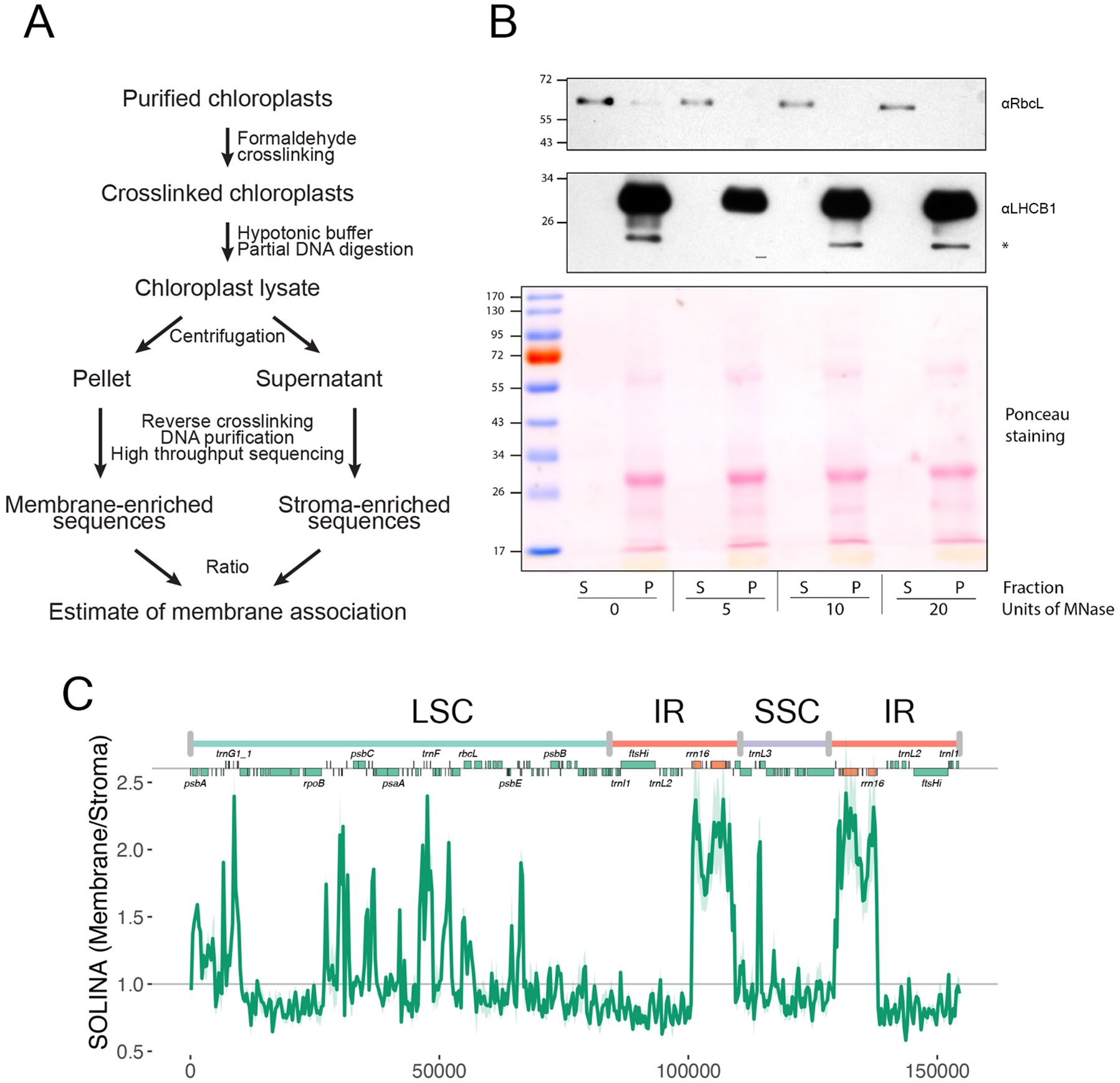
Chloroplast nucleoid is organized by association with the membranes A. Workflow of the <u>Sol</u>uble-<u>In</u>soluble Nucleoid Fractionation <u>A</u>ssay (SOLINA). B. Validation of SOLINA by western blot demonstrating presence of RbcL in the soluble (stroma) fraction and LHCB1 in the insoluble (membrane) fraction. Units of MNase and fractions are labelled on the bottom of the panel. S indicates soluble and P indicates pellet. Star indicates a non-specific band. C. Genome-wide pattern of membrane association identified by SOLINA. Signal from SOLINA in Col-0 wild-type plants was calculated in 50-bp genomic bins and plotted throughout the entire plastid genome. Genome annotation including genomic regions, positions of annotated genes ^22^, and names of selected individual genes is provided on top of the plot. Average enrichments from three independent biological replicates are shown. The light green ribbon indicates standard deviation.

To test the specificity of the SOLINA assay, we performed western blots with antibodies against membrane-and stroma-localized proteins. RbcL, the stroma-localized large subunit of Rubisco, was detectable only in the soluble fraction (Fig. 3B). In contrast, LHCB1, the membrane-localized subunit of the light-harvesting complex II, was detectable only in the insoluble fraction (Fig. 3B). This is consistent with previous observations ^32^ and confirms that our approach separates membrane and stromal fractions. To further confirm the specificity of SOLINA, we used the results of a prior study, which identified the region around 16S and 23S rRNA genes as membrane-bound ^27^. The outcome of SOLINA-seq was highly consistent with this observation (Fig. 3C), which further supports the specificity of our assay. While we cannot entirely exclude the possibility that properties other than membrane binding may affect DNA fractionation, we interpret the enrichment in insoluble fraction as evidence of membrane association.

Chloroplast DNA showed a complex sequence-specific pattern of membrane association in SOLINA (Fig. 3C). In addition to rRNA genes in the inverted repeats (IR), several other genomic regions in both the large single copy (LSC) and small single copy (SSC) were preferentially associated with the membranes (Fig. 3C). This indicates that preferential membrane association involves a subset of genes in a sequence-specific pattern. This suggests that the chloroplast nucleoid is organized by DNA anchoring to the membranes.

### Membrane association is correlated with PEP transcription

Preferential membrane association of rRNA genes and other genomic regions that contain highly transcribed genes (Fig. 3C) suggests that membrane association may be correlated with transcription. To test this hypothesis on the genome-wide scale, we compared membrane association determined by SOLINA with PEP binding to DNA determined by ptChIP-seq ^22^. Membrane association and PEP binding were strongly and significantly correlated on annotated genes (Fig. 4A) and throughout the entire plastid genome (Fig. S3A). Consistently, the strongest membrane association was observed on rRNA and tRNA genes (Fig. 4A) and was enriched on gene promoters (Fig. S3B), where PEP binding is also enriched ^22^.

**Figure 4.**
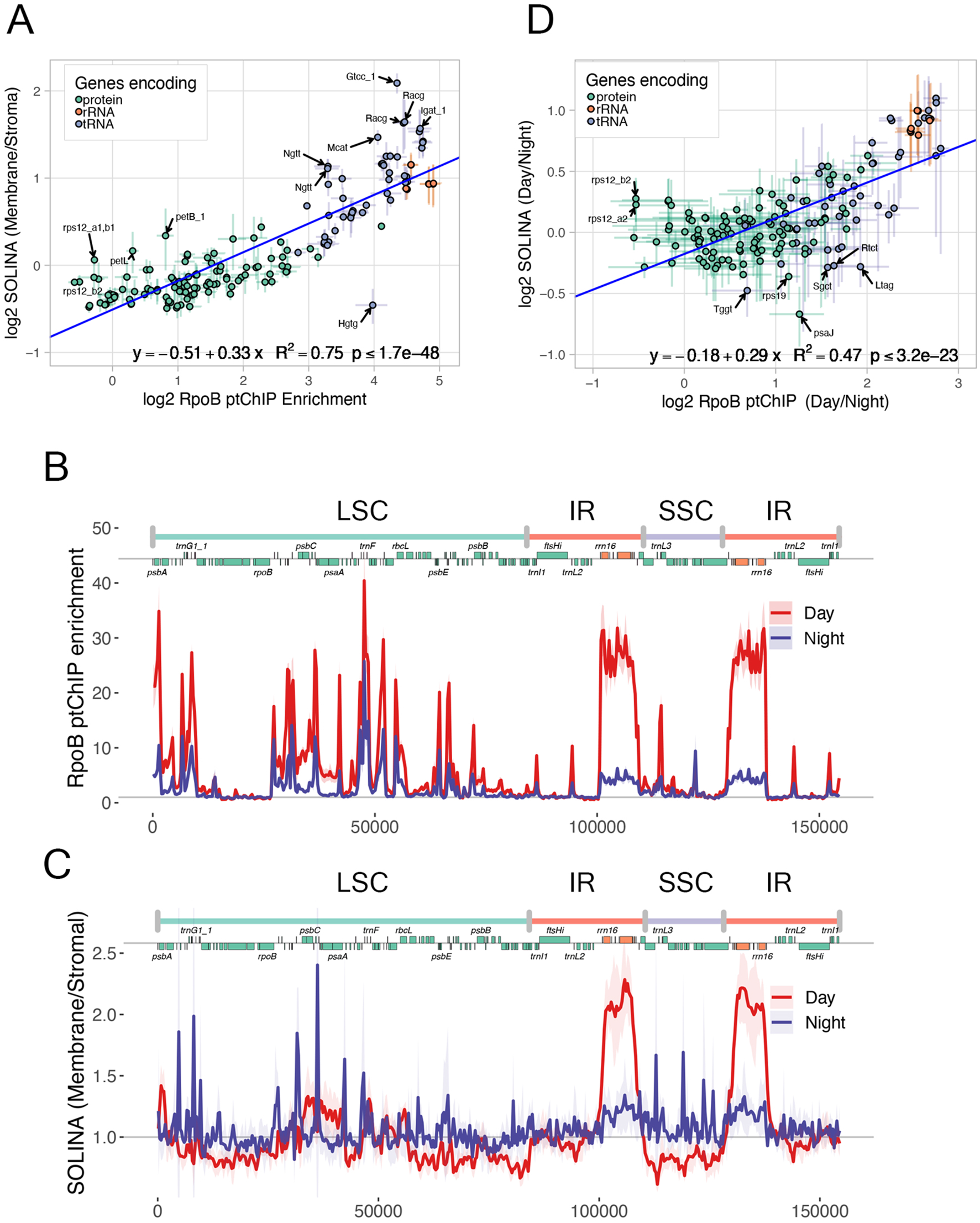
Membrane association is correlated with PEP transcription. A. Membrane association and PEP binding are significantly correlated. SOLINA signal and previously published RpoB ptChIP-seq signal ^22^ were compared on annotated genes. Data points are color-coded by function and show averages from three biological replicates. Error bars indicate standard deviations. The blue line represents the linear regression model. B. Extended dark treatment affects the pattern of PEP binding to DNA. RpoB ptChIP-seq was performed on Col-0 wild-type plants collected during the day or after extended dark treatment and enrichment was calculated in 50-bp genomic bins and plotted throughout the entire plastid genome. Genome annotation including genomic regions, positions of annotated genes ^22^, and names of selected individual genes is provided on top of the plot. Average enrichments from three independent biological replicates are shown. Ribbons indicate standard deviations. C. Extended dark treatment affects the pattern of membrane association. SOLINA was performed on Col-0 wild-type plants collected during the day or after extended dark treatment and signal was calculated in 50-bp genomic bins and plotted throughout the entire plastid genome. Average enrichments from three independent biological replicates are shown. Ribbons indicates standard deviations. D. Changes in membrane association and PEP binding after extended dark treatment are significantly correlated. Changes in SOLINA signal and RpoB ptChIP-seq signal were compared on annotated genes between plants collected during the day and after extended dark treatment. Data points are color-coded by function and show averages from three biological replicates. Error bars indicate standard deviations. The blue line represents the linear regression model.

To further validate the observed correlation between PEP binding and membrane association, we asked if the correlation persists in plants grown under different physiological conditions. Prolonged dark treatment is expected to result in a substantial change in the pattern of chloroplast gene expression ^32^. RpoB ptChIP-seq confirmed that 24-hour dark treatment leads to a genome-wide change in the pattern of PEP binding to DNA (Fig. 4B, Fig. S3C). Similarly, the pattern of DNA membrane association detected by SOLINA was also changed upon 24-hour dark treatment (Fig. 4C). Interestingly, the change in PEP binding to DNA was significantly correlated with the change in membrane association (Fig. 4D, Fig. S3D). This further confirms that membrane association is correlated with PEP binding to DNA.

Together, these results suggest that PEP-transcribed regions of the chloroplast genome are preferentially associated with the membranes. In contrast, non-transcribed sequences are not efficiently crosslinked to the membranes due to physical distance and/or lack of crosslinkable protein-DNA interactions.

### Membrane binding is correlated with local DNA-DNA interactions

Preferential membrane association of PEP-transcribed genomic regions suggests that binding to the membranes may be a general organizing principle of the chloroplast nucleoid. To determine the extent of this phenomenon, we adopted chromosome conformation capture ^33^ for use with fractionated plastid nucleoids. In this method, which we refer to as SOLINA-Hi-C, we performed crosslinking with EGS followed by crosslinking with formaldehyde. The first crosslinking step allowed for long-range protein-protein crosslinking to increase the sensitivity of the assay. This was followed by purification of the insoluble fraction like in SOLINA and identification of long-range chromosomal interactions by a modified Hi-C protocol.

SOLINA-Hi-C revealed no detectable long-range DNA-DNA interactions in the chloroplast nucleoid. This is shown by the lack of elevated signal away from the diagonal in Fig 5A. This observation indicates that at longer distances, chloroplast DNA is organized without sequence-specific and crosslinkable DNA-DNA interactions. Instead, long-range DNA organization is more likely to be stochastic.

**Figure 5.**
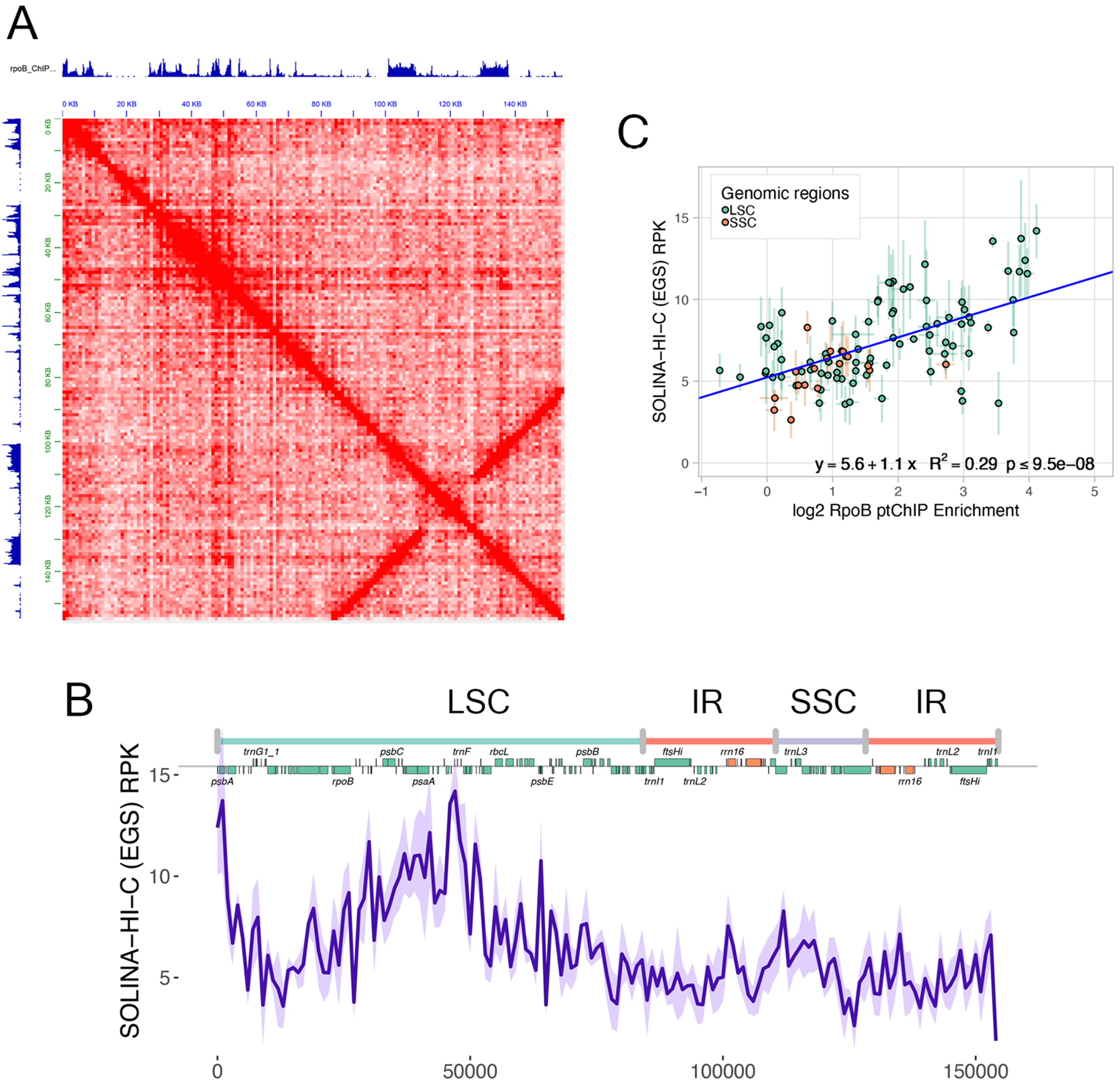
Membrane binding is correlated with local DNA-DNA interactions. A. Lack of long-range chromosomal interactions and presence of local interactions in the membrane-bound fraction of the chloroplast nucleoid. SOLINA-Hi-C data are plotted as a matrix in 1 kb bins. Strong signal across the diagonal is expected in the absence of any interactions. Counter-diagonal corresponds to inverted repeats. Previously published RpoB ptChIP-seq ^22^ is plotted on top and on the left. B. Presence of local DNA-DNA interactions in the chloroplast nucleoid. Frequency of interactions ranging between 3 kb and 10 kb is plotted in 1 kb bins. C. Local interactions in the membrane fraction and PEP binding are significantly correlated. SOLINA-Hi-C signal in the membrane fraction and previously published RpoB ptChIP-seq signal ^22^ were compared on 1 kb genomic bins within LSC and SSC regions. Data points are color-coded by region and show averages from three (RpoB ptChIP-seq) or four (SOLINA-Hi-C) biological replicates. Error bars indicate standard deviations. The blue line represents the linear regression model.

Further analysis of SOLINA-Hi-C results indicated the presence of local DNA-DNA interactions, which may be seen as signal close to the diagonal (Fig. 5A). They are most prominent in the 3kb to 10kb range, where a complex pattern of interactions may be observed (Fig. 5B). The frequency of short-range interactions was correlated with PEP binding in the LSC and SSC (Fig. 5C). Interestingly, this correlation did not apply to rRNA genes (IR regions), which despite being highly transcribed, showed low frequency of short-range interactions (Fig. S4A). These results indicate that highly transcribed genes in the LSC and SSC form local interactions at the membranes, but untranscribed genes adopt a more stochastic 3D organization.

### PEP transcription drives membrane association

The observed correlation between PEP binding to DNA and membrane association does not imply causality. To test if transcription is causing membrane association, we used mutants defective in sigma factors SIG2 and SIG6, which are known to directly recruit PEP to specific genes ^34–36^ and affect genome-wide patterns of PEP binding to DNA ^22^. SOLINA performed with *sig2* and *sig6* mutants revealed broad disruptions of DNA membrane association (Fig. 6A). Comparison of SOLINA to previously published RpoB ptChIP-seq in *sig2* ^22^ demonstrated a significant correlation between changes in PEP binding to DNA and changes in DNA association with the membranes (Fig. 6B). An even stronger correlation was observed in *sig6* (Fig. 6C), which has a more pronounced impact on PEP binding to DNA than *sig2* ^22^. This indicates that reduction of PEP binding to specific genes in sigma factor mutants leads to a reduction in membrane association at those genes. We interpret this result as evidence that transcription is a causal factor influencing the pattern of DNA-membrane association.

**Figure 6.**
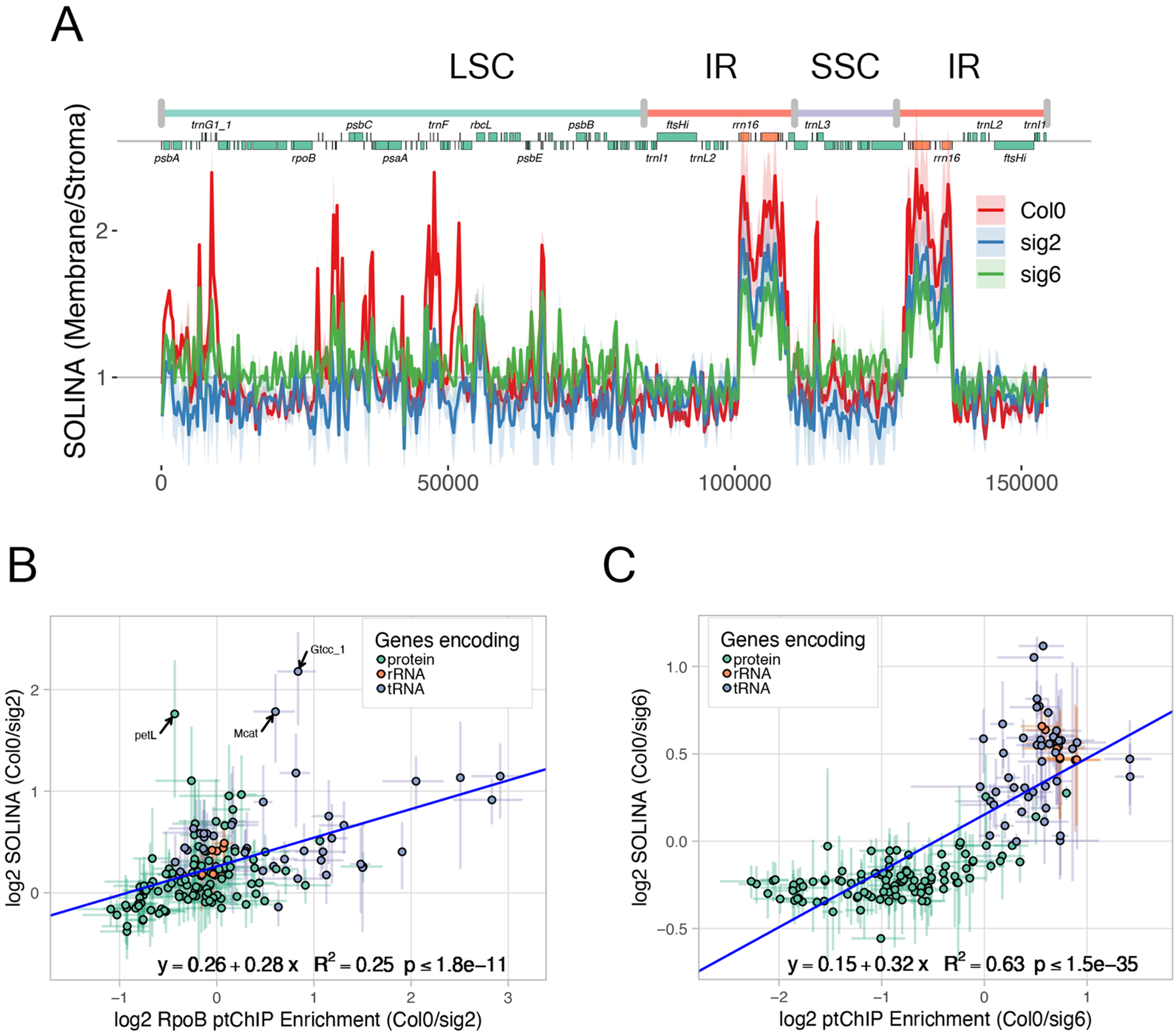
PEP transcription drives membrane association. A. Genome-wide pattern of membrane association in sigma factor mutants. Signal from SOLINA in Col-0 wild-type, *sig2* and *sig6* plants was calculated in 50-bp genomic bins and plotted throughout the entire plastid genome. Genome annotation including genomic regions, positions of annotated genes ^22^, and names of selected individual genes is provided on top of the plot. Average signal from three independent biological replicates is shown. Ribbons indicate standard deviations. B. Changes in membrane association are correlated with changes in PEP binding in the *sig2* mutant. Changes in SOLINA signal and previously published RpoB ptChIP-seq signal ^22^ between Col-0 wild-type and *sig2* mutant were compared on annotated genes. Data points are color-coded by function and show averages from three biological replicates. Error bars indicate standard deviations. The blue line represents the linear regression model. C. Changes in membrane association are correlated with changes in PEP binding in the *sig6* mutant. Changes in SOLINA signal and previously published RpoB ptChIP-seq signal ^22^ between Col-0 wild-type and *sig6* mutant were compared on annotated genes. Data points are color-coded by function and show averages from three biological replicates. Error bars indicate standard deviations. The blue line represents the linear regression model.

## DISCUSSION

We propose a model of chloroplast nucleoid organization where DNA is packaged in a sequence-specific manner and the main determinant of this specificity is the level of transcription. Highly transcribed genes like *psbA*, *rbcL*, many tRNA genes, and ribosomal RNA genes are preferentially attached to the thylakoid membranes. Membrane-associated DNA has high levels of overall protein occupancy, which is dominated by PEP. Other proteins may also bind to membrane-associated DNA, but this binding is either less prominent, weaker, less sequence-specific, or less susceptible to formaldehyde crosslinking. High protein binding instigates a certain level of DNA protection, likely caused by the widespread presence of PEP. Membrane association of highly transcribed genes leads to local 3D interactions within the nucleoid. Given their short distance, these interactions most likely occur only within individual transcriptional units, which indicates that each gene is recruited to the membrane independently. Transcription is causal in recruiting active genes to the membranes.

Regions of the genome with low levels of transcription are depleted in binding to the membranes. This may be caused by their presence in the stroma or by less efficient formaldehyde crosslinking to membrane proteins. DNA with low levels of transcription has low protein occupancy, no detectable DNA protection, and fewer local 3D interactions. This may be caused by an overall low amount or low sequence-specificity of protein binding or limited formaldehyde crosslinking of bound proteins.

Highly transcribed and membrane-associated DNA in our model likely corresponds to the biochemically detected Transcriptionally Active Chromosome and the central body observed using EM ^11, 15^. Untranscribed and unprotected DNA that is not associated with the membranes is consistent with the DNA fibrils observed in EM ^11, 15^. It should however be noted that there may be only limited equivalency of structures detected using *in vivo* and *in vitro* approaches. Moreover, our data only show enrichment of highly transcribed genes at the membranes and do not exclude the possibility of transcription occurring in the stroma.

Our model is based on the interpretation that the insoluble chloroplast fraction corresponds to the membranes and membrane-bound factors. This interpretation has strong support in the literature ^28–32^, protein composition of soluble and insoluble fractions, and consistency of SOLINA results with published data. It remains possible that properties other than direct binding to the membranes may drive certain molecules to the pellet during chloroplast fractionation, which is a potential limitation of our interpretation.

The role of transcription in controlling membrane association of DNA is most strongly supported by the observed impacts of sigma factor mutants. Although *sig2* and *sig6* affect transcription and also partially disrupt chloroplast membrane structure ^35, 36^, extended dark treatment affects transcription levels without impacting the membrane organization. Together, these results support the causal role of transcription in organizing the chloroplast genome. The impact of other processes like NEP transcription, replication, or DNA repair remains unknown. The causal relationship between transcription and DNA packaging indicates that the simplistic nuclear concept of heterochromatin as inaccessible and repressive does not apply to chloroplast nucleoids. Instead, transcription leads to more tight packaging and DNA inaccessibility to Tn5 transposase. Although our data put transcription upstream of DNA packaging, DNA organization may still play a role in achieving the proper pattern of gene expression.

The overall internal structure of the nucleoid does not involve complex 3D organization, like long-range looping or territories enriched in certain genomic regions or groups of genes, that could be detected by Hi-C. Instead, chloroplast DNA only interacts locally and likely adopts a more random arrangement at longer distances. Preferential membrane binding of highly transcribed genes is the most prominent detectable non-random property. However, Hi-C detects only interactions that are preserved by crosslinking ^37^, so weak protein-DNA interactions may lead to underestimation of long-range chromosomal interactions. Additionally, each chloroplast contains multiple nucleoids and each nucleoid contains multiple copies of the genome ^38^. Therefore, structural heterogeneity may also lead to long range interactions being underestimated.

Mechanisms that recruit transcribed DNA to the membranes remain unknown. Certain nucleoid-associated proteins like MFP1, PEND, TCP34 and pTAC16 are expected to directly bind to the membranes ^39–42^. If they also preferentially bind to transcribed DNA, they could contribute to DNA recruitment to the membranes. An alternative possibility is direct or indirect membrane binding of PEP ^32^. Yet another explanation is coupled transcription-translation-membrane insertion, also known as transertion ^43, 44^, which is however less likely, since chloroplast RNA tends to be highly stable and preferential membrane binding is also observed on genes encoding soluble proteins. Resolving the mechanism of membrane recruitment remains an important goal for future studies.

## EXPERIMENTAL PROCEDURES

### Plant materials and growth conditions

We used *Arabidopsis thaliana* wild-type Columbia-0 (Col-0) ecotype plants for all the experiments in this work, and we included the following genotypes: *sig2-2* (SALK_045706) (Woodson et al., 2012), and *sig6-1* (SAIL_893_C09) ^35^. Seeds were stratified in darkness at 4°C for 48 hours and grown on soil at 22°C under white LED light (100 μmol m^-2^ s^-1^) in 16h/8h day/night cycle for 14 days, or grown on 0.5X MS plates (0.215% MS salts, 0.05% MES-KOH pH 5.7, 0.65% agar) for 4 days at 22°C under constant white LED light (50 μmol m^-2^ s^-1^). For experiments requiring extended dark treatment, 14 days-old plants were exposed to 24-hour darkness treatment. Subsequent chloroplast isolation and cross-linking were performed avoiding light exposition. Control samples were collected after 3 hours of light exposition.

### Chloroplast enrichment and crosslinking

Chloroplasts from 14-day-old seedlings were enriched and crosslinked following the protocol adapted by ^22^ based on the original protocol from ^45^. In the case of SOLINA-Hi-C experiments, chloroplasts were subjected to protein-protein crosslinking with 5 mM EGS [ethylene-glycol bis(succinimidyl succinate))] solubilized in DMSO (dimethyl sulfoxide), for 45 minutes at room temperature and then subjected to formaldehyde crosslinking as reported by ^22^.

### Chloroplast IPOD

Protein occupancy in the chloroplast nucleoid was assayed by adapting the previously described IPOD technique ^20, 21^. Briefly, 50 – 100 µg crosslinked enriched chloroplasts [quantified by the amount of chlorophyll ^46^] were solubilized in 1X MNase reaction buffer (50 mM Tris-HCl pH 8.0, 5 mM CaCl_2_)] supplemented with 100 µg of RNase A and incubated for 20 minutes on ice; MNase was added to obtain fragments ranging from 100 to 200 bp and the sample was incubated for 10 minutes at 30 °C. MNase reaction was stopped by adding EDTA to a final concentration of 100 mM. The stopped reaction was combined with 1 volume of 100 mM Tris base and 1 µl of 10 % BSA. One volume of 25:24:1 phenol:chloroform:isoamyl alcohol was added to the sample, mixed, incubated for 10 minutes at room temperature, mixed again, and centrifuged at 21,130 g for 2 minutes at room temperature. The aqueous and organic phases were taken out and discarded by carefully bringing the interphase protein disk against the tube wall. The disk was resuspended by the addition of 1 volume TE buffer (10 mM Tris pH 8.0, 1 mM EDTA pH 8.0) and 1 volume Tris base, and extracted using 1 volume of 24:1 chloroform:isoamyl alcohol, mixed and centrifuged as before. The disk was resuspended in 1 volume of TE buffer and further isolated with 1 volume 24:1 chloroform:isoamyl as described. The final protein disk was solubilized in ChIP elution buffer (100 mM Tris pH 8.0, 10 mM EDTA, 1 % SDS) and subjected to reverse cross-linking and DNA isolation as reported by ^22, 47^.

### ptChIP-seq

ptChIP-seq experiments to detect RpoB binding to DNA were performed as described by ^22^.

### ptATAC-seq

DNA accessibility in the chloroplast nucleoid was assessed with Tn5 transposition by adapting a previously described nuclear ATAC-seq protocol ^26^. Briefly, one microgram of crosslinked enriched chloroplasts was resuspended in 1X Tn5 reaction buffer (Illumina) and assayed according to the manufacturer’s instructions. The transposed DNA was purified using the MinElute PCR purification kit following the manufacturer’s instructions. High throughput sequencing libraries were generated as reported by ^26^.

### SOLINA

Membrane and stromal-enriched DNA regions were identified by the Soluble-Insoluble Nucleoid Fractionation Assay (SOLINA). 100 µg of crosslinked enriched chloroplasts were resuspended in 1X MNase reaction buffer and incubated on ice for 20 minutes. MNase was added at different concentrations as shown in the results considering that shorter fragments (between 100 – 300 bp in the soluble fraction) increase the assay resolution, and incubated at 30°C for 10 minutes. MNase reaction was stopped by adding EDTA and EGTA at a final concentration of 10 mM each, and the sample was centrifuged at 21,130 g for 10 minutes at 4°C. The soluble fraction was transferred to a new tube, and the pellet was resuspended in 1 volume of ChIP elution buffer. SDS was added to the soluble fraction to reach a final concentration of 1%. Reverse crosslinking and DNA isolation for both fractions was performed as described by ^22, 47^.

### SOLINA-HI-C

To test for DNA-DNA contacts we based our approach on a previously described chromatin conformation capture protocol for pellet and supernatant ^33, 48^. Briefly, 100 µg of dual crosslinked enriched chloroplasts were resuspended in 1X DpnII reaction buffer (NEB) and digested with 20 U DpnII at 37°C overnight with gentle shaking. DpnII was stopped by heating the sample at 80°C for 20 minutes. The sample was centrifuged as in SOLINA protocol, and the pellet fraction was resuspended in 1X DpnII buffer. Ligation was performed by increasing the volume 5 times with T4 DNA ligase buffer (NEB) to a final concentration of 1X, adding 20 U of T4 DNA ligase, and incubating the sample overnight at 16°C with rotation. SDS was added to the ligated pellet fraction to reach a final volume of 1%. Reverse crosslinking and DNA isolation was performed as described by ^22, 47^.

### Immunoblot analysis

Proteins from SOLINA assays were extracted by resuspending samples in protein extraction buffer (20 mM Tris-HCl (pH 6.8), 3% β-mercaptoethanol, 2.5% sodium dodecyl sulfate, 10% sucrose) and incubated for 1.5 h at room temperature. Twenty micrograms of proteins from the pellet fraction and its equivalent volume of the supernatant fraction were separated by SDS-PAGE. To detect RbcL, polyclonal anti-RbcL antibody (PhytoAB catalog number PHY0096A) and anti-rabbit IgG antibody conjugated with horseradish peroxidase (Thermo catalog number PI314) were used as the primary and secondary antibodies, respectively. For LHCB1 detection we used the polyclonal anti-LHCB1 antibody from Agrisera (catalog number AS01 004) and the same secondary antibody; both detections were made at the same time by splitting the membranes in half considering the apparent protein molecular weights. Protein bands were visualized using chemiluminescence reagents (ECL Prime Western Blotting Detection Reagent, Amersham) and an ImageQuant LAS 4000 imager or blue films (Kodak).

### Data analysis

For chloroplast IPOD, ptChIP-seq, ptATAC-seq, and SOLINA, we performed the analysis as described by ^22^. For ptATAC-seq, genome-wide significant differences from 11 biological replicates were detected using the NBPseq ^49^. In the case of SOLINA-HI-C, the raw sequencing reads were trimmed using trim_galore v.0.4.1 and analyzed using Juicer ^50^. Contact matrices were drawn by merging the 4 biological replicates generated, and the matrices for each replicate were extracted by using juicer-tools ^50^. One-dimensional contact plots were generated by counting the normalized number of contacts from 3 to 10 kb distances for each 1 kb bin across the genome. The same data was used for the regression analyses.

## ACCESSION NUMBERS

The sequencing data from this study have been submitted to the NCBI Gene Expression Omnibus (GEO; http://www.ncbi.nlm.nih.gov/geo/) under accession number GSE228230. Sequencing data presented in this study are available through a dedicated publicly available Plastid Genome Visualization Tool (Plavisto) at http://plavisto.mcdb.lsa.umich.edu.

## ACKNOWLEDGEMENTS

This work was supported by a grant from the National Science Foundation (MCB 1934703) to A.T.W. S.F. was supported by grants from the Japanese Society for the Promotion of Science (19J01779, 20K15819). We thank Peter Freddolino for critical reading of the manuscript.

---

**Figure S1.**
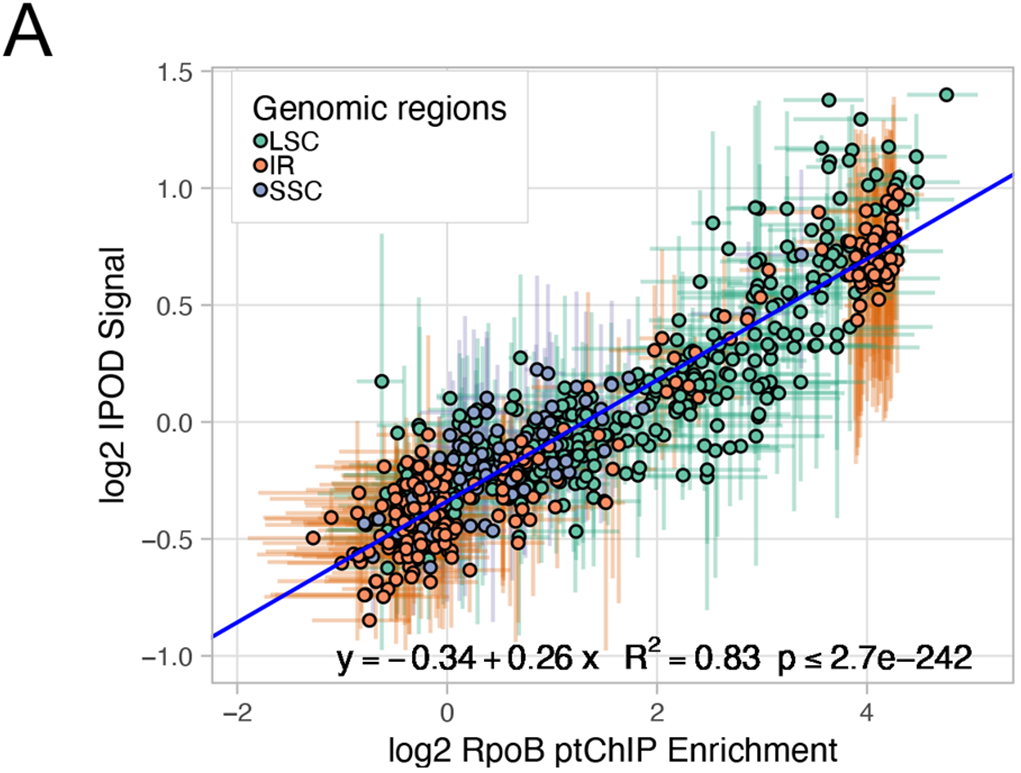
Protein binding to chloroplast DNA is dominated by PEP. A. Protein occupancy and PEP binding are significantly correlated. IPOD signal and previously published RpoB ptChIP-seq signal ^22^ were compared on 50 bp genomic bins. Data points are color-coded by genomic region and show averages from three (ptChIP-seq) or four (IPOD) biological replicates. Error bars indicate standard deviations. The blue line represents the linear regression model.

**Figure S2.**
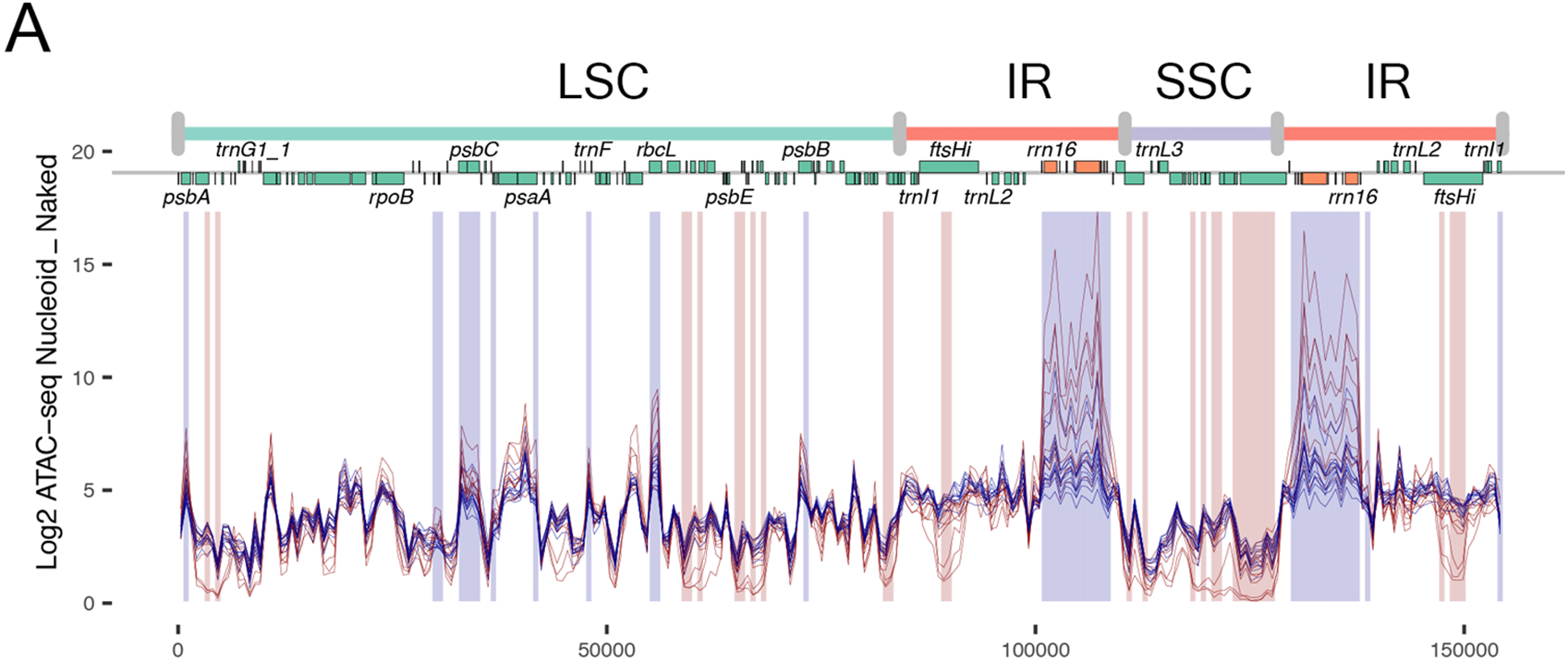
Transcribed genes have reduced DNA accessibility to Tn5 transposase. A. Individual biological replicates of ptATAC-seq data. Signal from ptATAC-seq in Col-0 wild-type plants was calculated in 50-bp genomic bins and plotted throughout the entire plastid genome. Red lines correspond to naked DNA, blue lines correspond to nucleoid DNA. Y axis represents normalized number of insertions. Genome annotation including genomic regions, positions of annotated genes ^22^, and names of selected individual genes is provided on top of the plot. Red shading indicates significant accessibility and blue shading indicates significant protection identified using a negative binomial model FDR < 0.05.

**Figure S3.**
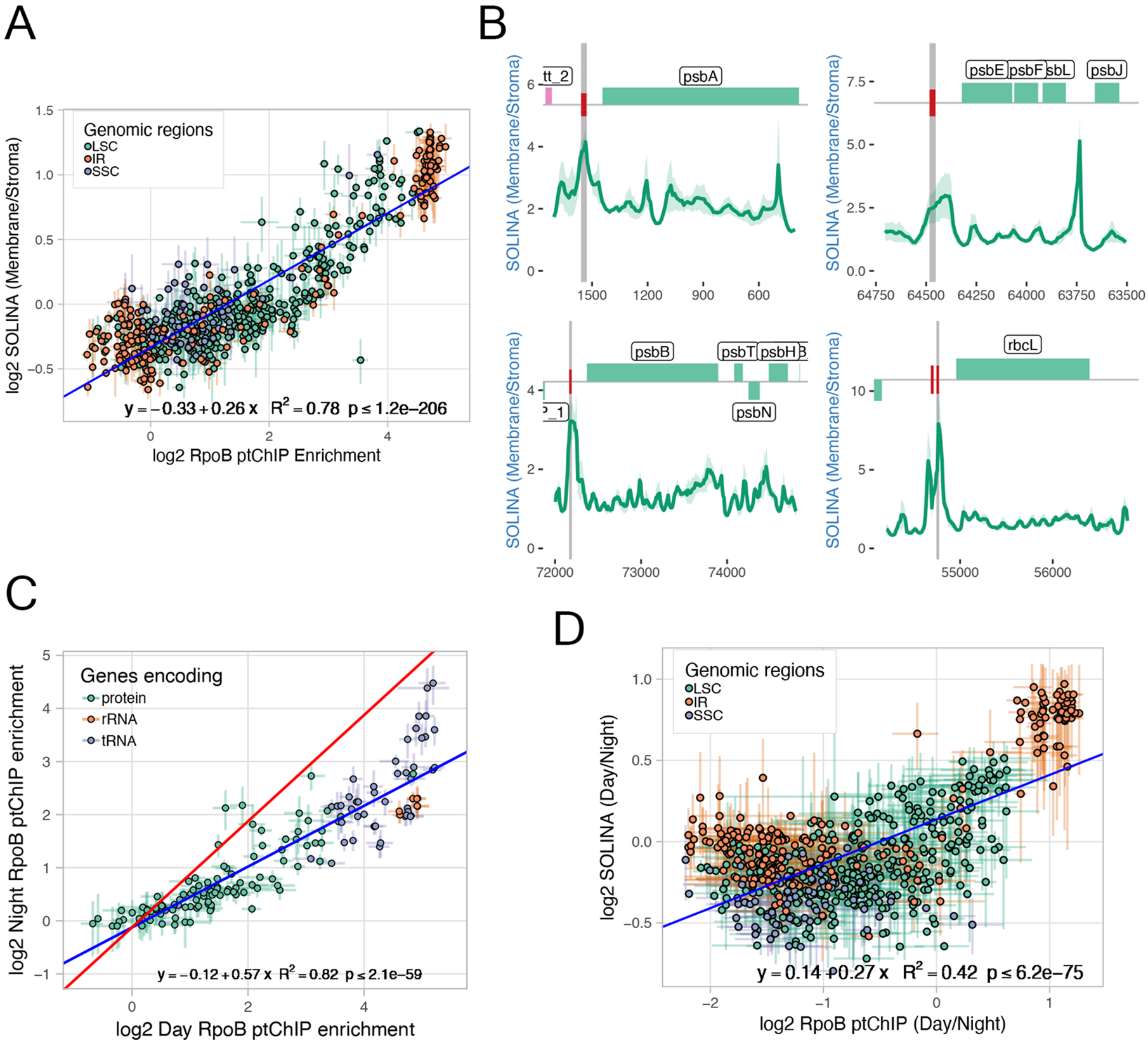
Membrane association is correlated with PEP transcription. A. Membrane association and PEP binding are significantly correlated. SOLINA signal and previously published RpoB ptChIP-seq signal ^22^ were compared on 50 bp genomic bins. Data points are color-coded by region and show averages from three biological replicates. Error bars indicate standard deviations. The blue line represents the linear regression model. B. Preferential membrane association on gene promoters. SOLINA signal enrichment from Col-0 wild-type plants was calculated in 10-bp genomic bins and plotted at *psbA*, *psbE*, *psbB* and *rbcL* loci. Average signal from three independent biological replicates is shown. Light green ribbons indicate standard deviations. Gray vertical lines indicate positions of the annotated promoters. Genome annotations are shown on top. C. Extended dark treatment affects the pattern of PEP binding to DNA. RpoB ptChIP-seq was performed on Col-0 wild-type plants collected during the day or after extended dark treatment, enrichment was calculated on annotated genes and compared between light and dark conditions. Data points are color-coded by function and show averages from three biological replicates. Error bars indicate standard deviations. The blue line represents the linear regression model. D. Membrane association and PEP binding changes during light and dark treatments are significantly correlated. SOLINA signal and RpoB ptChIP-seq signal after light and dark treatments were compared on 50 bp genomic bins. Data points are color-coded by region and show averages from three biological replicates. Error bars indicate standard deviations. The blue line represents the linear regression model.

**Figure S4.**
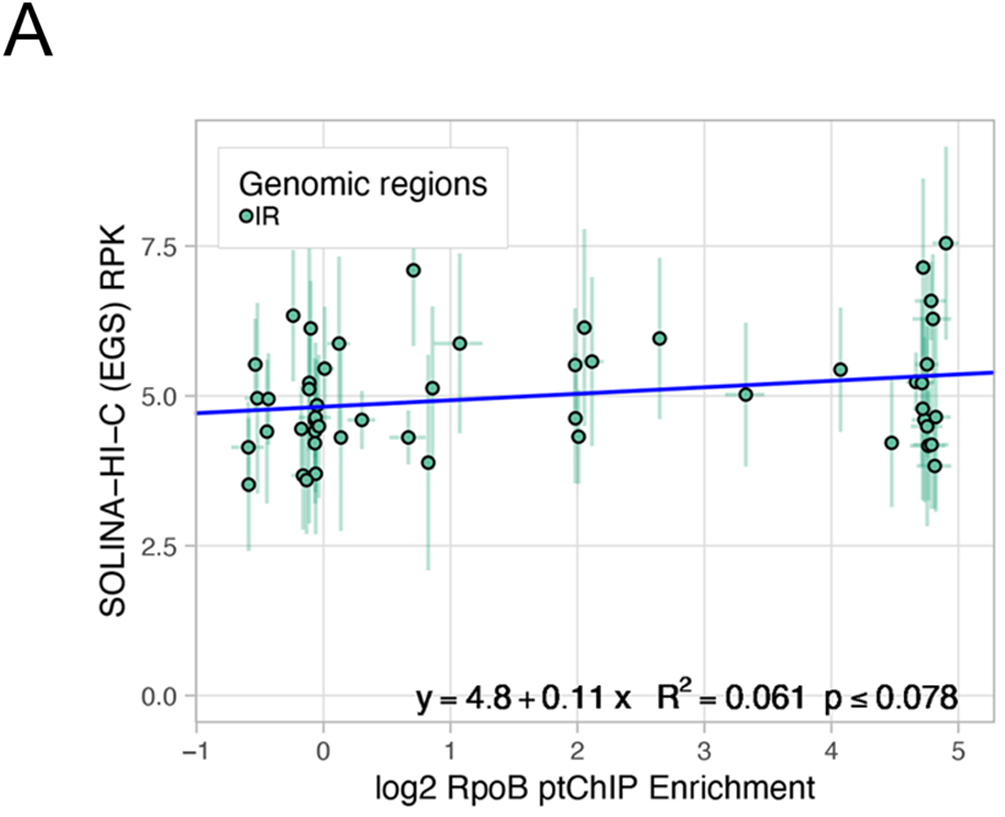
Membrane binding is correlated with local DNA-DNA interactions. A. Local interactions in the membrane fraction and PEP binding are not significantly correlated within the IR region. SOLINA-Hi-C signal in the membrane fraction and previously published RpoB ptChIP-seq signal ^22^ were compared on 1 kb genomic bins within the IR region. Data points are color-coded by region and show averages from three (RpoB ptChIP-seq) or four (SOLINA-Hi-C) biological replicates. Error bars indicate standard deviations. The blue line represents the linear regression model.

